# Mesoscopic oblique plane microscopy (Meso-OPM) with a diffractive light sheet- enabling large-scale 4D cellular resolution imaging

**DOI:** 10.1101/2022.03.29.486239

**Authors:** Wenjun Shao, Minzi Chang, Kevin Emmerich, Patrick O Kanold, Jeff S Mumm, Ji Yi

**Affiliations:** Department of Biomedical Engineering, Johns Hopkins University, Baltimore, Maryland, 21231, USA; Department of Ophthalmology, Johns Hopkins University, Baltimore, Maryland, 21231, USA

## Abstract

Fundamental understanding of large-scale dynamic connectivity within a living organism requires volumetric imaging over a large field of view (FOV) at biologically relevant speed and resolution. However, most microscopy methods make trade-offs between FOV and depth resolution, making it challenging to observe highly dynamic processes at cellular resolution in 3D across mesoscopic scales (e.g., whole zebrafish larva). To overcome this limitation, we have developed mesoscopic oblique plane microscopy (Meso-OPM) with a diffractive light sheet. By augmenting the illumination angle of the light sheet with a transmission grating, the axial resolution was improved ~6-fold over existing methods and ~2-fold beyond the diffraction limitation of the primary objective lens. We demonstrated an unprecedented FOV up to 5.4 × 3.3 mm with resolution of 2.5× 3 × 6 μm, allowing volumetric imaging of 3D cellular structures with a single scan. Applying Meso-OPM for *in vivo* imaging of zebrafish larvae, we report here the first *in toto* whole body volumetric recordings of neuronal activity at 2 Hz volume rate and the first example of whole body volumetric recordings of blood flow dynamics at 5 Hz with 3D cellular resolution.

## 1. Introduction

Modern microscopy, in conjunction with genetically-encoded fluorescent reporters, [1–4] has greatly advanced fundamental studies in neuroscience, cardiovascular biology, and developmental biology. As many dynamic biological processes are interconnected across anatomical structures, rapid high-resolution volumetric imaging over a wide FOV is critical to interrogating complex molecular and cellular interactions in living organisms[5–7]. For example, neuronal signals often traverse large distances within the brain to coordinate functions[6,7]. However, volumetric recording of neuronal activity patterns with sufficient FOV, resolution is a significant challenge due to the physical limit of optical diffraction. Microscopy techniques using a single primary objective are generally constrained by this limit, such as confocal, multiphoton, dictated by the numerical aperture (NA) of the objective used. This necessitates trade-offs between FOV and spatial resolution. For example, near-centimeter or multi-millimeter FOV can be obtained with sub-cellular lateral resolution by using low numerical aperture (NA) objective lens (*e.g*, NA = 0.35), but the axial resolution is typically above ~15 μm which is insufficient to resolve cellular features in the z-dimension [8–13]. The diffraction limit also extends to microscopy methods that deliver illumination and collect fluorescence via the same refractive optics. For example, a computational miniature mesoscope used a micro-lens array instead of a regular objective lens to overcome the trade-off between FOV and resolution. Yet, the axial resolution is limited by the total angular coverage of the micro-lens array leading to axial resolution of tens of microns[14].

The diffraction limit also poses an additional challenge in that axial resolution deteriorates more rapidly than lateral resolution with increasing FOV, since the depth of focus is inversely proportional to the square of NA. Thus, a much higher NA is needed to achieve axial cellular resolution than cellular lateral resolution. As a reference point, NA of ~0.5-0.6 are needed for two-photon (900 nm excitation wavelength) and single-photon (488 nm excitation wavelength) excitation to achieve cellular axial resolution, respectively[15,16]. Whereas the FOV of a high NA objective lens is fundamentally confounded by the scale-dependent geometric aberrations of its optical elements[17], the increase in NA is at the cost of FOV. Consequently, the FOV is usually less than 1 mm^2^ [18–23] when utilizing high NA objective lens, which is insufficient for imaging dynamic events over large scales. While tremendous efforts have been made to design objectives to achieve large-scale recording with cellular resolution, particularly in the z-dimension, the physical limit of diffraction still persists[9,12,15,16], and the complexity of the optics and imaging system also increases dramatically.

Another strategy for overcoming the diffraction limit is to introduce additional optical elements so that resolution is not solely defined by the primary objectives. The prominent example is to use two objectives to decouple excitation and detection as in light sheet microscopy (LSM)[24,25], selective plane illumination microscopy[26], dual-inverted selective-plane illumination microscopy[27], lattice light-sheet microscopy[28,29], light-sheet theta microscopy [30], and open-top light-sheet microscopes[31]. While the primary objective offers lateral resolution, the excitation lens independently creates a thin optical light sheet therefore the axial resolution is no longer constrained by the primary objectives.

Because of the diffraction limit, mesoscale volumetric imaging of live samples over multi-millimeter scales with 3D cellular resolution remains a tremendous challenge. In this paper, we introduce Meso-OPM which employs a novel strategy of a diffractive light sheet to overcome the inherent limitations mentioned above. Meso-OPM belongs to a family generally termed single objective light sheet microscopy (SOLSM) [32,33]. In contrast to conventional LSM, which uses two objective lenses, SOLSM uses only a single primary objective lens but applies an off-axis oblique light sheet excitation at the specimen and a remote focusing system to capture the scanning light sheet. The advantage of SOLSM is the system simplicity in that much of the optics is shared for both excitation and collection, as well as flexible sample mounting as in conventional upright or inverted microscope setting. Because regular SOLSM uses one primary objective, the resolution limitations described above still apply. Our previous Meso-OPM version achieved 5×6 mm^2^ FOV, but depth resolution is sacrificed to ~35 μm [13]. Here, the reported Meso-OPM addressed this persistent problem by creating an high angle illumination light sheet with a transmission grating. This enables the axial resolution to be improved by 6-fold, i.e., to cellular level even using a low NA primary objective (e.g, NA = 0.3). With 3D cellular resolution, we demonstrate an unprecedented FOV of 5.4 × 3.3 × 0.33 mm in an acute brain slice preparation at a resolution of 2.5× 3 × 6 μm, to resolve individual neurons. Using Meso-OPM to image living larval zebrafish, we also demonstrated whole-brain cellular resolution imaging of calcium dynamics at 2 Hz and whole-body blood flow imaging at 5 Hz.

## 2. Materials and methods

### 2.1 The working principle of Meso-OPM

The limitation of previous mesoscopic OPM methods is the inherent trade-off between FOV and resolution, particularly in depth direction[10,13]. For low NA objective (e.g., NA=0.3), the depth resolution is on the order of tens of microns. To circumvent this trade-off, we introduced a diffractive light sheet by using a transmission grating (Fig. 1a). Large angle diffractive light sheet can be created in 1^st^ order while maintaining the full collection angle for emission collection in 0^th^ order. The diffraction by the grating is independent of the primary objective lens, and thus not constrained by the diffraction trade-off discussed above. The concept for the proposed Meso-OPM system is shown in Fig. 1b. The light sheet angle mainly relies on the diffraction angle such that the axial resolution is improved 6-fold [10,13] compared to our previous mesoscopic OPM without grating (Fig. 1c) and a 2-fold of the diffraction-limited axial resolution (Fig. S1 and S2), thus achieving 3D cellular resolution. We compared FOV and depth resolution by Meso-OPM with an array of SOLSM techniques[10,13,19,32–38] (Fig. 1d), and found that our method maintains multi-millimeter FOV as well as micron-level depth resolution. Consequently, Meso-OPM can afford capturing 4D biological dynamics at cellular resolution across FOV dimensions that have been challenging to achieve previously.

**Fig. 1.**
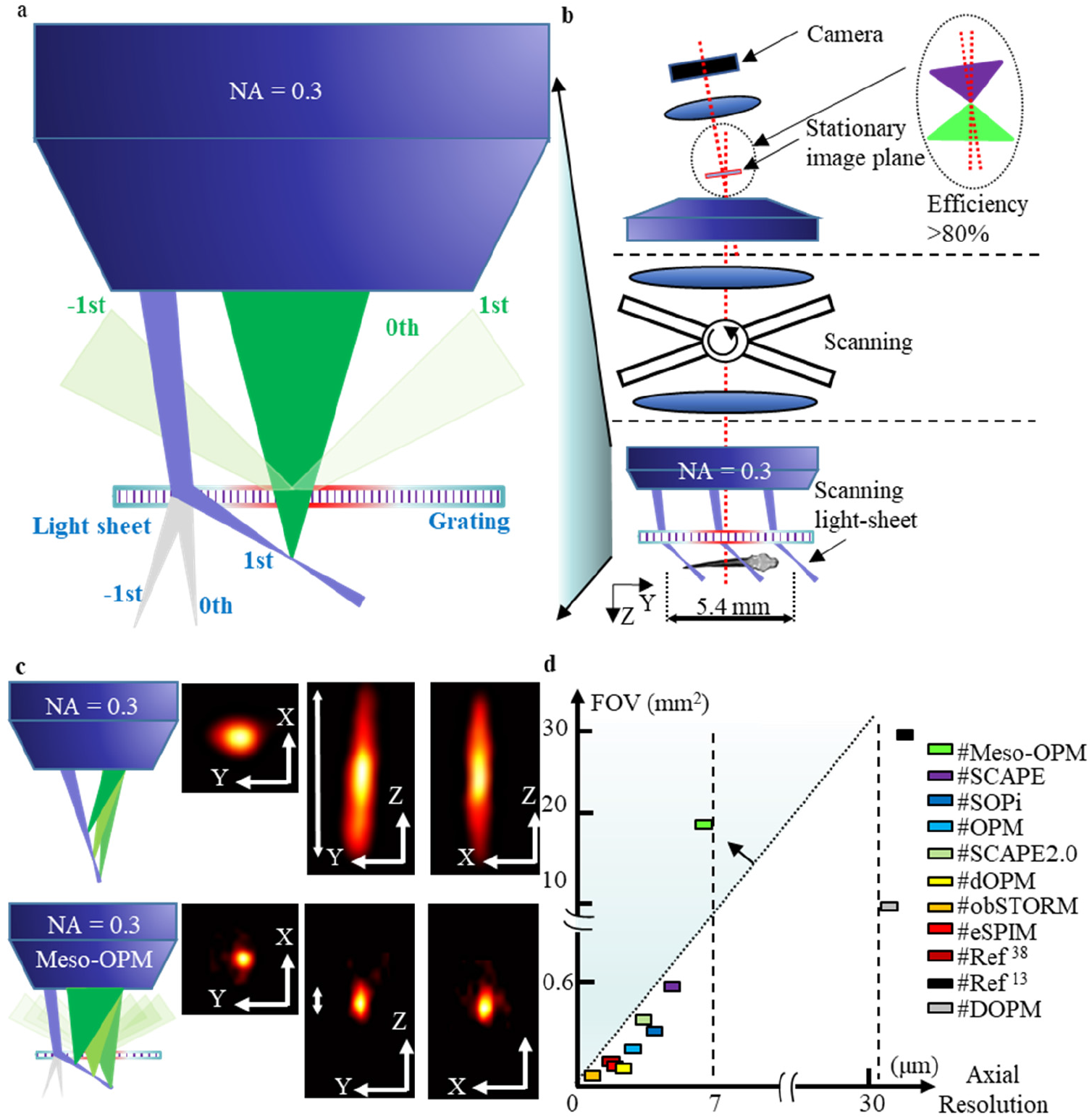
The concept of volumetric imaging with a diffractive light sheet. (**a**) High-angle light sheet illumination in 1^st^ order and full NA fluorescence collection in 0^th^ order with the help of transmission grating. (**b)** The concept of utilizing diffractive light sheet in Meso-OPM to achieve large FOV and 3D cellular resolution; (**c)** The comparison of spatial resolution with existing mesoscopic OPM. **(d)** The comparison of the FOV and axial resolution between different SOLSMs.

### 2.2 System description

The layout of the optical design for Meso-OPM is shown in Fig. 2a) (See Supplementary Fig. S3 for actual setup photograph). The key component that enables 3D cellular resolution over large FOV in Meso-OPM is the transmission grating (TG) shown in Fig. 2b). The excitation light sheet (blue) in Fig. 2b) can be diffracted into different orders by the flowing equation:

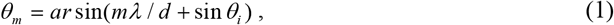

where *θ_m_* is the diffraction angle for *m*^th^ order, *m* represents the diffracted order, λ is the wavelength of the incident beam, *θ_i_* is the incident angle, *d* is the line density of the TG. We used the 1^st^ order diffraction for excitation, and collected the 0^th^ order fluorescence emission. Fluorescence in non-zero orders is either rejected by the aperture of the objective lens (OL1) or widely separated in the image space (Fig. 2c and Supplementary Fig. S4). With a larger illumination angle by the grating diffraction, the axial resolution is improved beyond the limitation set by the NA of the primary objective lens (See supplementary Fig. S6d and 6e). Having described the mechanism of creating and imaging with the high-angle diffractive light sheet, we set out for detailed explanations of the whole system below.

**Fig. 2.**
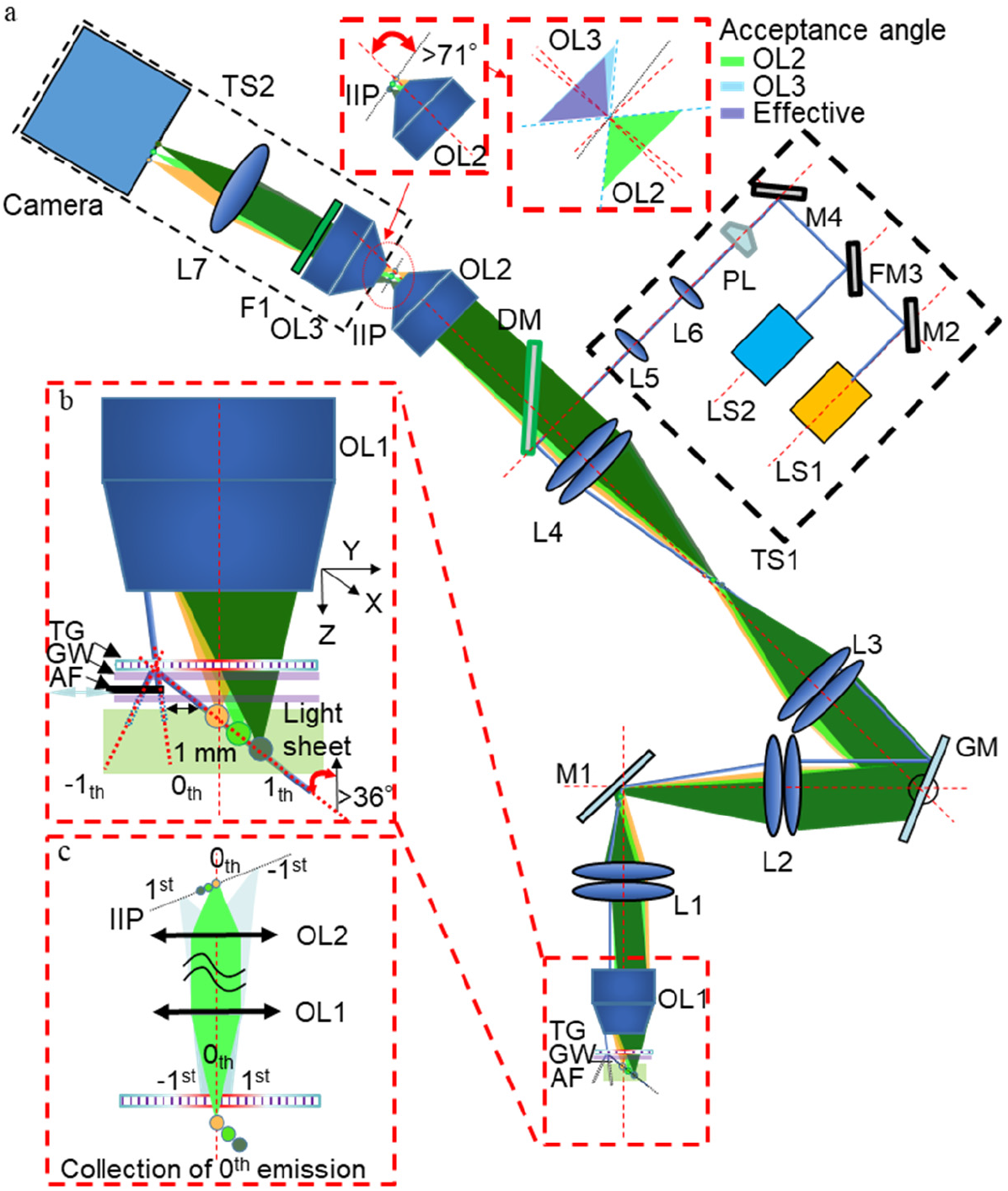
The schematic of the experiment setup. (**a**) The layout of the whole optical design. TG: transmission grating; GW: glass window; AF: aluminum foil; OL: objective lens; L: lens; M: mirror; GM: galvanometer mirror; IIP: intermediate image plane; F: filter; TS: translation stage; PL: Powell lens; FM: flip mirror; LS: light source. (**b**) The zoom-in view of the primary objective lens exhibiting the generation of high-angle excitation light sheet. The primary objective lens consists of the objective lens (OL1), transmission grating (TG), glass window (GW), the motorized aluminum foil (AF). (**c**) A simplified layout of the optics from the light sheet to the IIP showing the imaging of emission fluorescence in different diffraction orders.

The light source unit consists of two lasers (LS1: Coherent, 488nm, 50 mW; LS2: Coherent, 561nm, 100 mW), a Powell lens (PL: Edmund optics, 30° fan angle), a telescope (L5: Thorlabs, two AC254-250-A, *f_L5_* = 125 mm; L6: two AC254-050-A, *f_L6_* = 25 mm), two plane mirrors (M2 & M4: Thorlabs, PF10-03-P01), and one flip mirror (FM3: Thorlabs, FP90, and PF10-03-P01). Excitation light can be switched between LS1 and LS2 by FM3. PL has a fun angle of 30° and is used to create light sheet excitation. The light sheet was introduced into the microscope by a dichroic mirror (DM, Chroma, ZT488/561rpc-UF1). The light source unit is mounted on a translational stage to adjust the laser offset from the optical axis for oblique illumination.

The laser is then directed by two relay telescopes (L4, L3, L2: Thorlabs, two AC508-200-A, *f_L4_* = *f_L3_* = *f_L2_* = 100 mm; L1: Thorlabs, two AC508-100-A, *f_L1_* = 50 mm) and a scanning galvanometer mirror (Nutfield: QS-12 OPD, 20 mm aperture) to the back pupil of the primary objective (OL1, Olympus, UplanFL10×/0.3, *f_OL1_* = 18 mm). The offset of the beam at the back pupil of OL1 is ~3.5mm. As a result, the light sheet has an incident angle of ~11° when projected on the transmission grating (TG: Thorlabs, line density of 0.83 μm/line). The angle of the 1^st^ order diffraction in the air can be calculated by Eq. 1 to be ~51° and ~60° for 488nm and 561nm, respectively. The oblique angle *θ_r_* in water is then ~36° and ~41° correspondingly by refraction (See supplementary Fig. S5a). The diffraction efficiencies in 1^st^ order for 488nm and 561nm excitation are measured to be ~26% and ~20% while the diffraction efficiencies in 0^th^ order for the green and red emission are both ~ 35%.

Beyond the transmission grating, two coverslips were used to protect grating surface and for water immersion (Fig. 2b). The working distance from the OL1 assembly is ~1-2 mm. The distance between 0^th^, 1^st^ and −1^st^ excitation is ~1 mm. As long as the sample size is <1mm, no interference from 0^th^ and −1^st^ order excitation is present. For imaging large specimen, such as brain slice, the 0^th^ and −1^st^ orders of the diffraction were blocked by a moving aluminum foil (AF) sandwiched between two coverslips (Fig. 2b and supplementary Fig. S3b and 3c) to avoid undesired fluorescence excitation.

As for excitation wavelength of 488 nm, the beam waist (*ω_0_*) and the Rayleigh (*Z_R_*) range in the immersion water are approximately 5 μm and 214 μm, respectively. As for 561nm, the beam waist and Rayleigh range are slightly bigger, which are 6 μm and 267 μm, respectively (See the calculation with Eq. S2 and S3 in the supplement). The depth imaging range is then estimated by 2*Z_R_cosθ_r_*, where *θ_r_* is the oblique angle of the light sheet in water (*i.e*. 36° and 41 ° for 488 and 561 nm, see supplementary Fig. S5a), to be 346 and 403 μm.

The 0^th^ fluorescence emission through the grating is collected by the system. The light sheet image is directed by the relay lens L1 and L2 and de-scanned by GM. A stationary intermediate image plane (IIP) can be formed after OL2 (Olympus, UPLSAPO 20×/0.75, *f_OL2_* = 9 mm). The additional glass substrate of the transmission grating, as well as the immersion water, may increase spherical aberration. To mitigate spherical aberration, the magnification from L1 to L4 was designed to be 2 (*f_L2_×f_L4_/f_L1_/f_L3_* = 2) so that the back aperture of the OL2 is overfilled to reject high angle rays at the cost of light loss (See supplementary Eq. S6). According to the simulation in Fig. S8, the optical system from OL1 to OL2 is less sensitive to glass and immersion media induced aberration. The optical performance can be significantly improved by better optical design and customized transmission grating, which will be discussed in the end.

The lateral magnification from the specimen to IIP can be calculated according to the focal length of lenses as *M*=*f_OL2_× f_L1_×f_L3_/f_OL1_/f_L2_/f_L4_* =0.25. Given the magnification (*M* = 0.25) and the angle of the light sheet (*θ_r_* = 36° for 488 nm, 41° for 561 nm), the angle (*θ_IIP_*) of the light sheet image in IIP can be calculated as follows [13]:

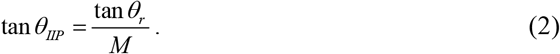

As for excitation wavelengths of 488 nm and 561 nm, *θ_IIP_* is calculated to be ~71° and 74° in air respectively. Finally, a remote focusing system, consisting of objective lens (OL3: Olympus, UPLFLN40×/0.75, *f_OL3_* = 4.5 mm), filter (F1: Chroma, MF525-39; Thorlabs, AT575lp), camera lens (L7: Navitar, MVL75M1, *f_L7_* = 75 mm), and camera (Andor, Zyla 4.2), is correspondingly rotated by 19° or 16° for collection. Because of the augmented light sheet angle by grating, the IIP is almost perpendicular to the optical axis (zoom-in of the IIP in Fig. 2a). Therefore, >80% of the light from OL2 to OL3 can be collected, resulting in an effective NA (lateral) of ~0.15 (See Fig. S7).

### 2.3 The theoretical resolution

After the description of the setup, we now give the theoretical expression of the system resolution. As the angle of the excitation light sheet is independent of the NA of the primary objective lens (OL1), the calculation of the system point spread function (PSF) can be written as follows:

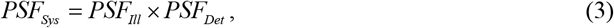

where *PSF_Ill_* is the illumination PSF, *PSF_Det_* is the detection PSF. The calculation is the same as that for the LSM except for the non-orthogonality between the illumination and detection PSF. A numerical simulation based on our previous publications[39,40] is used to calculate the theoretical resolution according to Eq.3 (See supplementary Fig. S6a). We first established 3D coherent amplitude transfer function, and then simulated PSFs for both illumination and emission. The calculated theoretical resolutions are ~1.4 μm (X) × 1.6 μm (Y) × 6 μm (Z) for 488 nm excitation and ~1.5 μm (X) × 1.8 μm (Y) × 6 μm (Z) for 561nm excitation. The axial resolution, as well as the depth imaging range under varying illumination angles resulting from different line densities of the TG, are analyzed (See supplementary Fig. S6f).

### 2.4 Data processing

As a result of the oblique illumination light sheet, the acquired image volume is a stack of oblique 2D images. Therefore, affine transformation consisting of both shearing and scaling is applied to the volume data to reconstruct the actual geometry of the sample [10,34,39].

As for Fig. 5b and supplement video 4, the lower part of the fish data (mainly the digestive system) is removed to highlight the neuronal features in the brain and the spinal cord. To generate the 2D angiogram shown in Fig. 6d-g, every two adjacent volume data sets in the whole time series are subtracted to firstly generate a differential time series. Then, a 3D angiogram of the entire fish can be created by applying MIP along the time axis of the differential time series. Lastly, the projection (Fig. 6d-g) was obtained by taking MIP along each dimension (X, Y, and Z) of the 3D angiogram.

### 2.5 Biological sample preparation

All animal-related procedures were in accordance with the Institutional Animal Care and Use Committee at Johns Hopkins University and conformed to the guidelines on the Use of Animals from the National Institutes of Health (NIH).

Red and green fluorescent microspheres (Polysciences: YG Microspheres, 1.00μm; Red Dyed Microsphere, 1.00μm) were used in the experiments for resolution characterization. They were diluted and immobilized in 1% agarose and molded in two separate Petri dishes. The Petri dish was covered with a cover glass (thickness: 0.1 mm, refractive index: ~1.52) to flat the gel surface. Water immersion is applied between the protective glass window of the objective lens and the cover glass before imaging.

Transgenic zebrafish larvae expressing green reef coral fluorescent protein in endothelial cells (Tg(VEGFR2:GRCFP)ZNL, Fig. 3), green fluorescent protein in macrophage cells (Tg(spi1b:GAL4,UAS:EGFP), Fig. 6), and genetically encoded calcium indicators (Tg(elavl3:jGCaMP7s), Fig. 5) were used in the *in vivo* experiments. Zebrafish at 4-5 dpf with length of 3-4 mm were cultivated at ~ 28°C following standard procedures (light cycle conditions of 14h light and 10h dark). To mount samples, zebrafish larvae were first anesthetized with tricaine (MS-222) for ~3 minutes and then added to heated 1.5% low-melting agarose in a Petri dish to allow for orientation at room temperature before providing overlayed E3 media following agarose hardening. The refractive index of the water is 1.33. The objective was immersed into the Petri dish to image the fish directly in an upright way. After the imaging, all of the zebrafish were released into freshwater for recovery.

**Fig. 3.**
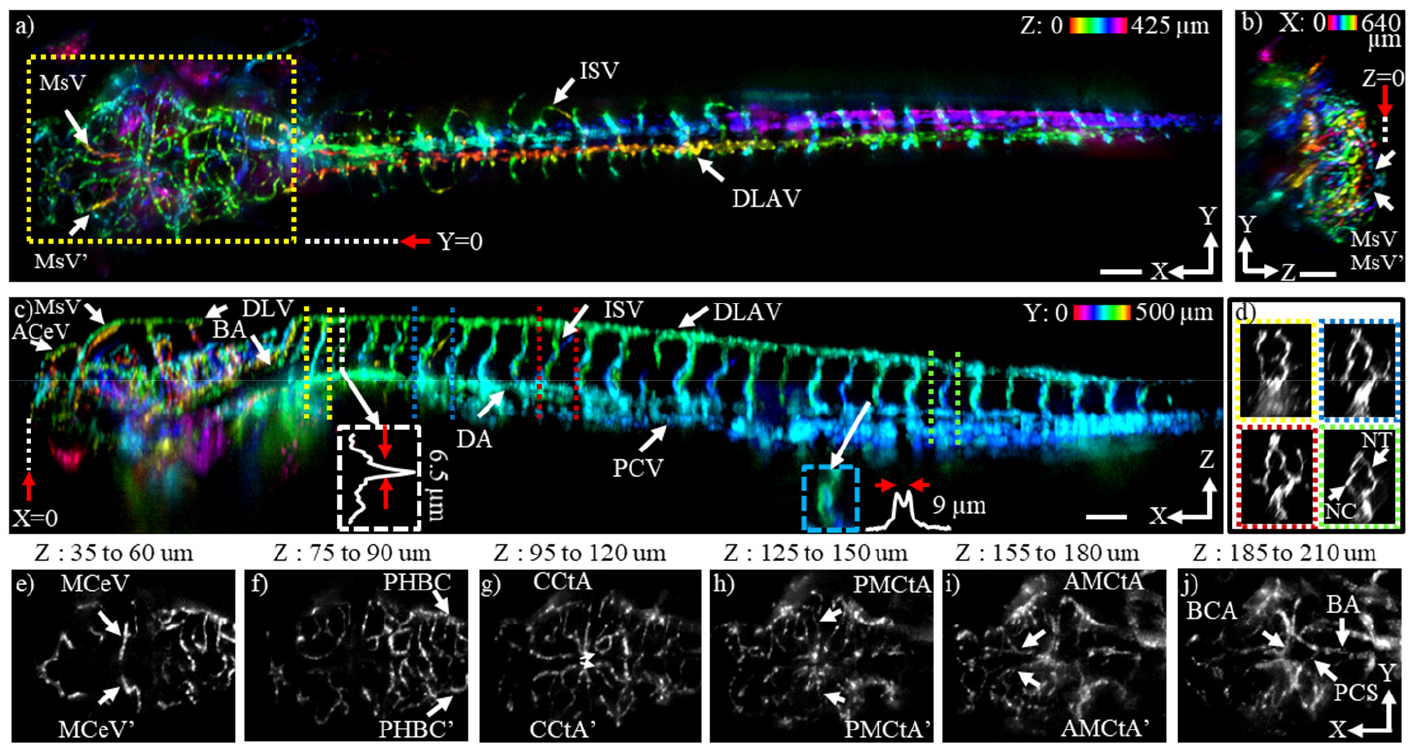
High-definition whole-body vasculature imaging of live zebrafish larva obtained in a single FOV at frame rate of 500 Hz. (**a**) Color-coded maximum intensity projection (MIP) in the XY plane over 425 μm along the Z direction. The 0 reference position of the MIP is marked by a red arrow in panel b; (**b**) Color-coded MIP in the YZ plane over 640 μm along the X direction indicated by the yellow dashed box in panel a. The 0 reference position of the MIP is marked by a red in panel c; (**c**) Color-coded MIP in XZ plane over 500 μm along the Y direction. The 0 reference position of the MIP is marked by a red arrow in panel a. (**d)** YZ cross-sections of 4 pairs of ISVs indicted by corresponding double dash lines in panel c; (**e-j)** *Enface* z-projections of the head region marked by yellow dashed box in panel a. Paired vessel structures such as MCeV (Middle cerebral vein), PHBC (Primordial hindbrain channel), CCtA (Cerebellar central artery), PMCtA (Posterior mesencephalic central artery), AMCtA (Anterior mesencephalic central artery), and PCS (Posterior communicating segment) can be seen within different projections. Scale bar, 100 μm. (BCA: Basal communicating artery; BA: Basilar artery.)

For the *in vitro* slice imaging, progeny from homozygous Calb2-IRES-cre mice (JAX no. #010774) crossed with homozygous tdtomato reporter mice (JAX no. #007909) was used. To prepare the acute brain slice, the whole brain of a 4-weeks-old mouse was dissected out in artificial cerebrospinal fluid ACSF containing (in mM): 130 NaCl, 3 KCl, 1.25 NaH2PO4, 20 NaHCO3, 10 glucose, 1.3 MgSO4.7H2O, and 2.5 CaCl2.2H2O (pH 7.35–7.4, equilibrated with 95% O2–5% CO2). A 400-μm tissue block containing both thalamus and cortex region (Cruikshank et al., 2002) was sliced in icy-cold oxygenated ACSF, using a microtome (Leica VT1200) and transferred into oxygenated ACSF warmed at physiological temperature (35–37°C) for an hour. After recovery, the acute brain slice was transferred to a perfusion chamber cycled with oxygenated ACSF at room temperature for imaging. The refractive index of the ACSF is 1.33. The objective lens was immersed into the perfusion chamber during the imaging.

## 3. Results

For each demonstration shown below, the imaging parameters are summarized in the supplementary material (Table S1) while the sample preparation is provided in Methods.

### 3.1 FOV and resolution characterization

As the angle of the diffractive light sheet is also a function of the excitation wavelength, we performed imaging experiments on two types of fluorescent microspheres to calibrate the FOV and resolution under blue and green light excitation. The lateral scale was calibrated by imaging the resolution target. To calibrate the the axial scale, the sample was moved in Z dimension by a manual positioning stage during imaging. The FOV under different excitation wavelengths is the same, which is 3.3 mm (X) × 5.4 mm (Y) × 0.33 mm (Z) (Supplement Fig. S1 and S2) obtained at 1 mm working distance. Because the illumination is augmented by the TG, the imaging range in Y is inversely proportional to the working distance, which can be adjusted to accommodate different applications (See supplementary Fig. S9). The resolution was quantified by calculating the full width at half maximum (FWHM) of the line intensity along X, Y, and Z directions of different beads. The resolutions along X, Y, and Z directions for blue light (488 nm laser) excitation are 2.5 ± 1.1 μm, 3 ± 1.4 μm, and 6 ± 1.8 μm, respectively, while that for green light (561 nm laser) excitation are 2.9 ± 1 μm, 3.5 ± 1.5 μm, and 6 ± 1.9 μm, respectively (Supplement Fig. S1 and S2). By dividing the accessible FOV of 3.3 mm × 5.4 mm× 0.33 mm by the volumetric resolution of 2.5 μm ×3 μm× 6 μm, the information throughput of our Meso-OPM can be estimated to be ~1.3 × 10^8^ resolvable image points across the imaged volume. By extension, this suggests the ability to monitor 4D signal dynamics of ~1.3 × 10^7^ cells.

### 3.2 High-resolution structure imaging in whole larval zebrafish and uncleared acute mouse brain slice

#### 3.2.1 Whole larval zebrafish imaging with Partial FOV

After characterizing the FOV and resolution, we carried out static structural imaging of large living samples to test system performance. We first imaged transgenic zebrafish larvae that express a green fluorescent protein reporter in vasculature endothelial cells at 4 days post fertilization (dpf). The approximate dimensions of a zebrafish larvae at this age are ~3-4 mm long, ~0.6 mm wide, and ~0.5 mm thick [41]. The laser power on the sample was ~ 3.4 mW. The acquired volume size has dimensions of 3.3 mm (X) × 0.65 mm (Y) × 0.55 mm (Z) with pixel density of 2048 × 300 × 275 pixels at a frame rate of ~500 Hz). The image quality is best within the calibrated FOV of ~0.33 mm in the Z dimension. By achieving 3D cellular resolution over a wider and deeper FOV, we can provide a high-definition 3D rendering of the whole larva (See supplement video 1).

Color-coded projections of volumetric data demonstrate the ability of Meso-OPM to resolve individual blood vessels throughout larval zebrafish in 3D (Fig. 3a-c). The branches of the mesencephalic vein (MsV) in the dorsal head region and the intersegmental veins (ISV) in the ventral trunk can be observed in both the lateral and axial projections (Fig. 3a-c). Thanks to the large FOV, the whole dorsal longitudinal anastomotic vessel (DLAV), which has a length of ~ 3 mm, can be fully visualized in the lateral (Fig. 3a) and axial views (Fig. 3c). The DLAV is connected to either dorsal aorta (DA) or posterior (caudal) cardinal vein (PCV) by ~29 pairs of ISVs over a distance of ~3 mm (Fig. 3c). The ability to discriminate all ISVs demonstrates that a depth penetration of ~250 μm can be maintained throughout the FOV (Fig. 3c). The intensity profile of the small feature along the depth direction suggests depth resolution better than 6.5 μm (white dash box shown in Fig. 3c), which is sufficient to resolve individual vessels in the depth direction. The double-edge structure of the ISV vessels can be revealed by the intensity profile of a single ISV (blue dash box shown in Fig. 3c). Transverse views of the trunk at four rostral-caudal positions clearly reveal the round shape of the neural tube (NT) and notochord (NC) (Fig. 3d) which typically have a diameter of ~25 μm and 50 μm, respectively[42]. *Enface* z-projections of the head region at six different depths demonstrate the ability to resolve individual vessels throughout the imaging volume (Fig. 3e-j). The discrimination of different vessel patterns, such as the paired brain vascular structures, indicates the depth resolution and penetration can be well maintained in the peripheral area of the FOV.

#### 3.2.2 Uncleared acute mouse brain slice imaging with Full FOV

We next imaged an acute mouse brain slice with calretinin-neurons expressing tdTomato under the full imaging volume of 3.3 mm (X) × 5.4 mm (Y) × 0.33 mm (Z) with a pixel density of 2048 × 2000 × 160 pixels at frame rate of 100 Hz. The laser power on the sample was ~ 2 mW. To eliminate the background by the 0^th^ and −1^st^ order diffraction by the grating, a synchronized moving aluminum foil is used as a beam blocker allowing only 1^st^ order diffraction excitation during the acquisition (See Fig. S3d in supplementary material). Figure 4a shows a single scan color-coded mesoscopic FOV covering major anatomical structures such as cortex, hippocampus, and thalamus. The uniform somatic structures indicate the cellular resolution is well maintained over the full FOV. Neuronal somas are clearly visualized from cortical layer I to IV in all three projections (Fig. 4a-4d), confirming 3D cellular resolution. The large FOV allows us to simultaneously capture neurons in the hippocampus (Fig. 4e and 4f) and the thalamus (Fig. 4g and 4h). Close examination demonstrates the capability to image apical dendrites of pyramidal neurons in layer VI (Fig. 4i). A high-definition 3D rendering of the acquired volume can be found in supplementary video 2.

**Fig. 4.**
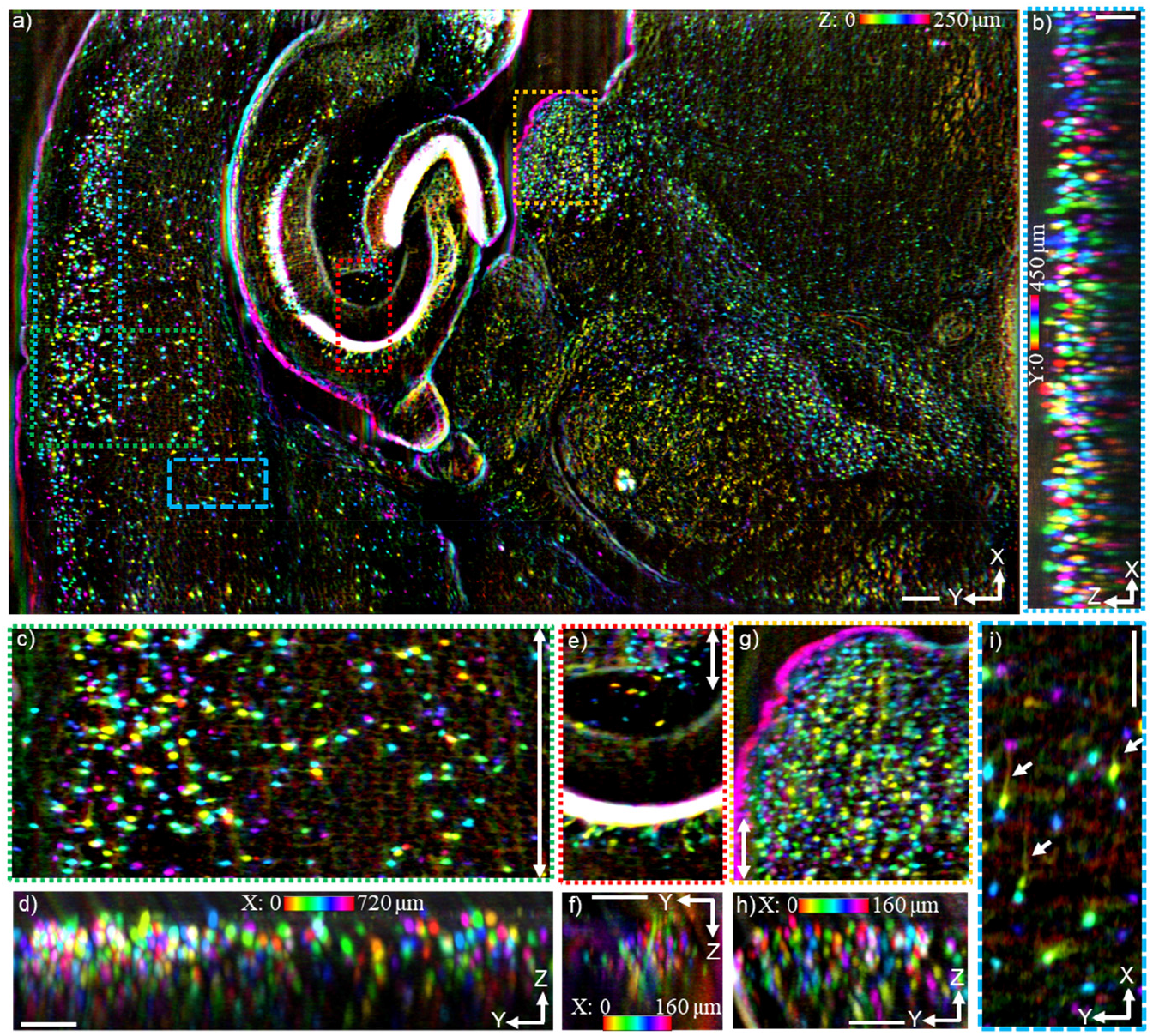
High-definition 3D imaging of an acute brain slice obtained in a single FOV at 3D cellular resolution across a wide FOV. (**a)** Color-coded MIPs of the *enface* view over a distance of 250 μm in the Z dimension, i.e., the whole brain slice volume. (**b**) Color-coded MIPs of the XZ plane over a distance of 450 μm in the Y dimension. The 0 reference positions of the projections are indicated by blue dash lines in panel a; (**c, e, and g**) Zoomed images of the *enface* views of the areas indicated by dashed boxes in panel a; (**d, f, and h)** Color-coded MIPs of the YZ plane of the volume indicated by double arrow lines in panel (c, e, and g). The scale bar for panel (b) is 200 μm; the scale bars for the other panels are all 100 μm.

**Fig. 5.**
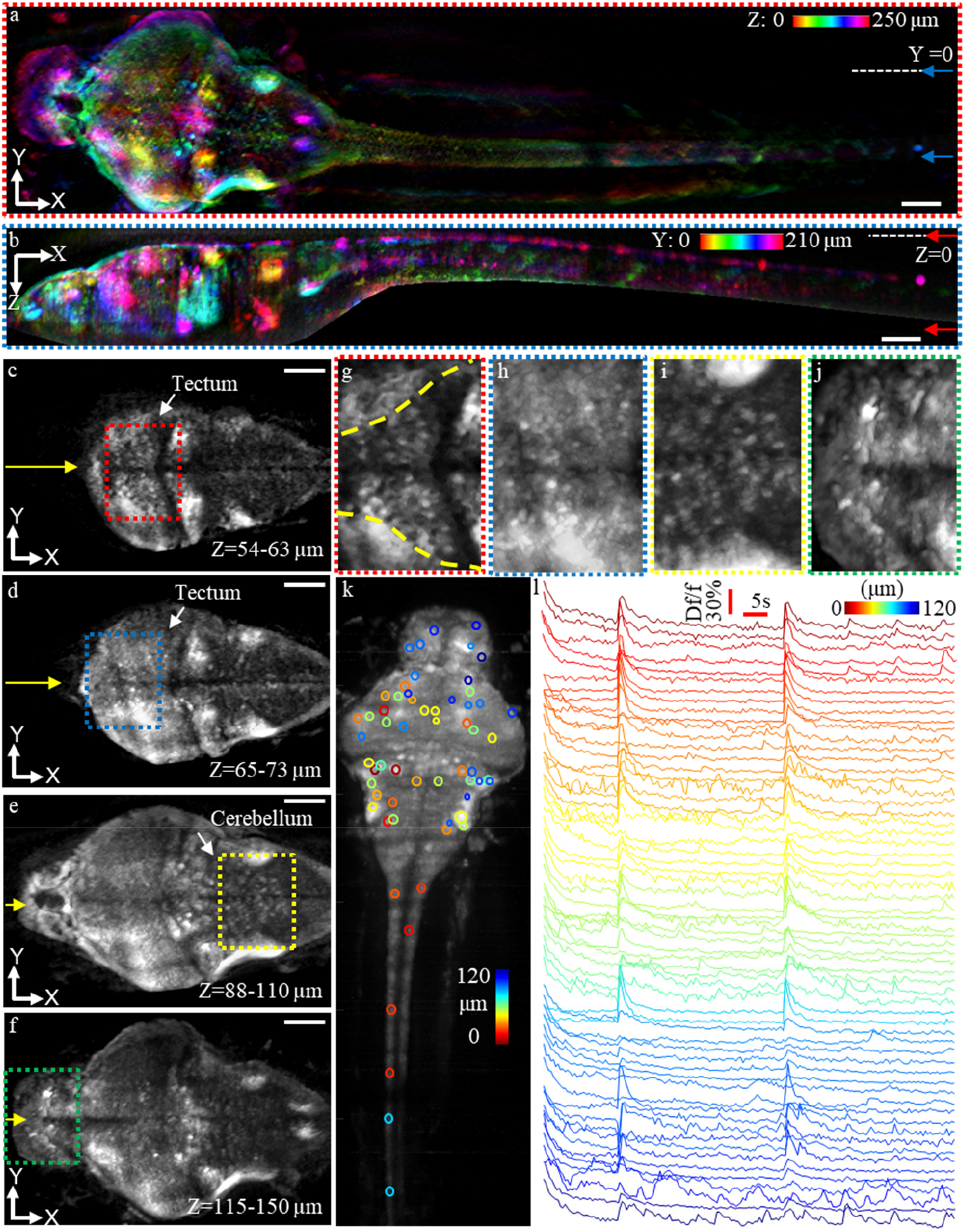
Whole-body neuronal activity recording in zebrafish larva with 3D cellular resolution. (**a**) Color-coded MIP in XY plane over 250 μm along the Z dimension. The reference positions of the MIP are marked by arrows in panel b. (**b**) Color-coded MIP in XZ plane over 210 μm in the Y dimension. The reference positions of the MIP are marked by arrows in panel a. (**c-f)** *Enface* projections at different z-depths across the whole brain. (**g-j)** Zoomed views of the boxed areas in panels c-f. **(k)** Representative neurons selected for evaluating correlated activity patterns are marked with circles, the color represents relative imaging depth from 0 to 120 μm. Fish data shown in a-k were acquired at 250 Hz per plane and scanned 125 planes at ~4 μm spacing, thus a 2 Hz whole-volume resolution. (**l**) Calcium traces of the selected neurons (see panel l) over an 85 s recording period. Scale bar, 100 μm.

**Fig. 6.**
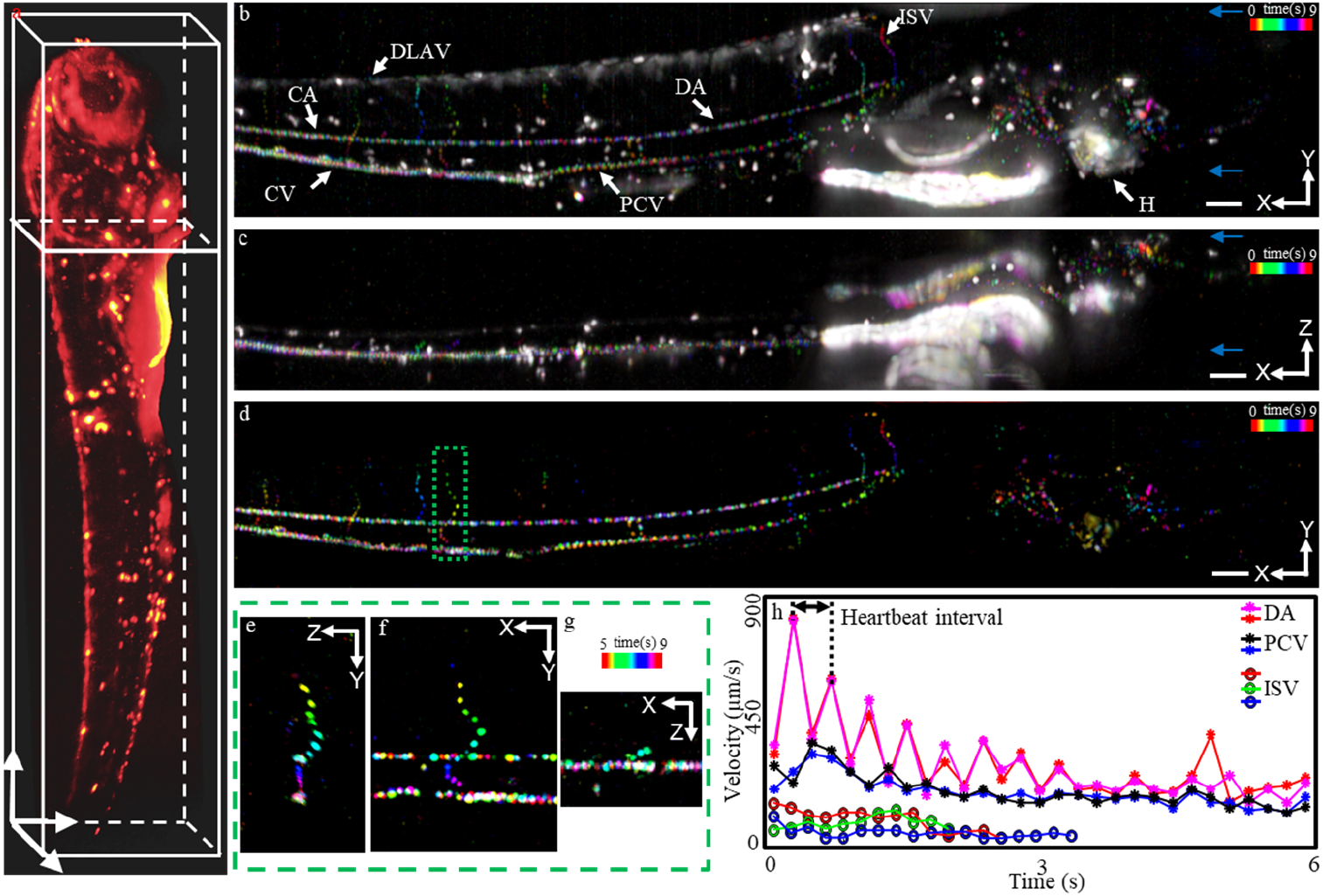
Imaging blood flow over the whole larval zebrafish at high spatiotemporal resolution. (**a**) Single volume captured at volume rate of 1 Hz to highlight the structure. (**b**) Color-coded MIP of cell movements in the XY plane. The XY view is generated by spatial MIP along Z direction over the position indicated by blue arrows in panel c; (**c**) Color-coded MIP of cell movements in the XZ plane. The XZ view is generated by spatial MIP along Y direction over the position indicated by blue arrows in panel b; (**d)** Color-coded angiogram in XY plane. (**e-g**) The tracking of blood cells in 3D over a time window of 4 s in a single ISV indicated in panel e. Fish data shown in b-g were acquired at 625 Hz per plane and scanning 125 planes at ~4 μm spacing, thus a 5 Hz whole-volume resolution. (**h)** Measurement of velocities of blood cells in different types of vessels. The heart rate was estimated to be ~140 Hz by Fourier analysis. Scale bar, 100 μm.

### 3.3 Whole-body recording of neuronal activity in zebrafish larvae

After characterizing FOV and resolution in static images of living samples, we proceeded to image neuronal signaling dynamics across an entire zebrafish larva expressing a cytosolic genetically encoded calcium indicator (jGCaMP7s) in neurons throughout the central nervous system. The acquired volume of the whole 4-dpf zebrafish has dimensions of ~ 3.3 mm × 0.5 mm × 0.4 mm. The laser power on the sample was ~ 3.4 mW with which no significant photon bleaching was observed during the imaging process. Color-coded maximum intensity projections (MIPs) of the whole zebrafish larvae are shown in Fig. 5a-c. The entire volume data has a pixel dimension of 2048 × 125 × 200 pixels (each volume contains 125 planes) at 2 Hz volume rate. Thanks to the large FOV, individual neuronal features can be identified throughout the entire nerve system from the brain to the spinal cord, as shown in the *enface* projection (Fig. 5a), as well as the side projection (Fig. 5b). MIPs of different brain regions at varying depths were generated (Fig. 5c-f). Distinct features can be clearly delineated, such as the boundary of the hemispheres indicated by the yellow arrows in Fig. 5c-f and the boundary of the optic tectum in Fig. 5g. For instance, tectal neurons in the midbrain can be clearly identified (Fig. 5c, g and 5d, h), as well as neurons in the cerebellum (Fig. 5e, i). Pallium and habenula neurons that are not present in Fig. 5c-e) can be seen in a deeper layer MIP (Fig. 5f, j), which agrees with the anatomy structure of the zebrafish brain[43,44]. Zoomed images of the boxed areas in Fig. 5g-j further confirm that individual neurons can be resolved at different tissue depths. A comparison of the image acquired with the Meso-OPM and the confocal microscope is provided in supplement material (Fig. S10) showing the similar anatomical organization of neurons.

Spontaneous calcium dynamics over the entire zebrafish nervous system were captured at a volume rate of 2 Hz for 85s. 60 neurons were manually selected at different depths throughout the brain and spinal cord (Fig. 5l) and fluorescence intensity traces plotted (Fig. 5n). These data show that at two time points many of the selected neurons appeared to fire synchronously. This correlated firing pattern can also be observed in supplementary video 3. As the FOV covers the whole larva, Meso-OPM offers a unique opportunity to study how neuronal activities are correlated over large volumes encompassing the entire brain and spinal cord, a level of inquiry that has been inaccessible to date. A 4D rendering of the time series can also be found in supplementary video 3.

### 3.4 Imaging blood flow over the whole larval zebrafish at high spatiotemporal resolution

By achieving large-field 3D cellular resolution across entire sample volumes without moving the sample or objective lens, our novel Meso-OPM method is well-suited for imaging cellular dynamics across whole organisms or whole tissues at fast speeds. As an additional demonstration of this, we performed volumetric imaging of blood flow using a transgenic zebrafish expressing enhanced green fluorescent protein (EGFP) in myeloid cells. The laser power on the sample is ~ 3.4 mW. The structure of the entire larval fish was first confirmed by the single volume shown in Fig. 6a. Next, images were acquired across a whole 4-dpf zebrafish larva (an ~3.3 mm × 0.5 mm × 0.4 mm volume) with pixel dimensions of 2048 × 125 × 200 pixels at a volume rate of 5 Hz (See supplement video 4). Results of timelapse imaging over a 9s time window show individual cell dynamics within major vessels of the circulatory system, such as the CA, caudal vein (CV), dorsal longitudinal anastomotic vessel (DLAV), intersegmental veins (ISV), posterior (caudal) cardinal vein (PCV), and dorsal aorta (DA) (Fig. 6b and c). Cell movements are indicated by color-coded temporal traces (Fig. 6b and c). The discrimination of individual blood cell movements in 3D across the whole volume, from heart to tail, further demonstrates the high temporal and spatial resolution possible with Meso-OPM (See 4D rendering in supplementary video 4). To quantify blood flow, motion contrast angiography (See *Material and methods*) was performed on the whole time series to highlight the motion and suppress the static signals as shown in Fig. 6d.

To exemplify the power of the high spatiotemporal resolution achieved, the 4D trajectory of a single cell traveling within an ISV from the DLAV to the CV over a time window of 4 s is provided in Fig. 6e-g. With the capability of tracking individual cells, we are able to quantify the speed of blood flow in different types of vessels such as the DA, PCV, ISV (Fig. 6h). The measured results agree with previous publications[45] revealing different velocities in each vessel type. For example, the velocity of the blood cell in the DA is faster than in the PCV or ISV as expected. Similarly, the pulsatile movement of the blood cells in the DA and PCV is more apparent than in ISV. Owing to the large FOV, blood cells in the DA were tracked over a distance of ~2 mm from the heart to the tail. An interesting observation is that the velocity in the DA gradually slows down as the blood cell are traveling away from the heart.

## 4. Discussion

Our diffractive mesoscopic OPM leverages the large FOV of off-the-shelf low magnification objective lens (e.g., 0.3 NA) while overcoming the limitation of insufficient axial resolution. The use of diffractive light sheet in Meso-OPM allows 3D cellular resolution over a 3.3 × 5.4 mm^2^ FOV without any mechanical translation. We demonstrated the performance by structural imaging of entire zebrafish larvae and a large acute brain slice. With large FOV and 3D cellular resolution, and biologically relevant speed, we were able to record calcium dynamics over the entire zebrafish nerve system at volume rate of 2 Hz, which was previously unattainable due to insufficient depth resolution[10,13] or limited FOV covering only the brain[18,25]. We also demonstrated single blood cell 4D tracking at volume rate of 5 Hz over the whole larval zebrafish. Imaging and quantifying dynamic cellular and signaling events between multiple tissues/regions, such as the recording of spontaneous neuronal activity between the brain and spinal cord, and blood cell circulation between multiple vessel types (e.g., from the DA to CA), has been an unmet challenge which Meso-OPM can overcome.

There are limitations of the current system that can be significantly improved to allow higher resolution and imaging speed. The light collection efficiency is sub-optimal in the current configuration. The off-the-shelve transmission grating has about 1/3 efficiency at 1^st^ and −1^st^ diffraction so that 0^th^ order fluorescence emission is limited. The current setup also overfilled the back pupil of OL2 causing light loss to mitigate the spherical aberration. To address this limitation, we can optimize the grating design to first have polarization selectivity such that >90% efficiency can be achieved in 1^st^ order diffraction for excitation wavelength while other high orders are negligible. Therefore, the motorized aluminum foil can be removed, and high volumetric imaging speed can be obtained over the full FOV rather than partial FOV. The better grating design would also allow the 0^th^ order efficiency to approach 50%. While this light loss is inevitable, the loss is compensated by high collection efficiency between OL2 and OL3 to maintain a reasonable overall light efficiency. The optical magnification can be also carefully designed to match the back pupils of OL1 and OL2 and avoid light loss. We can improve the aberration by using thinner grating, better relay optics, and objective lens with correction collar, as well as adaptive optics.

As for future work, we will explore primary objective lenses with different NA to explore combinations of FOV and resolution. Considering the diffraction efficiency can be optimized, the proposed method only requires dry objectives without any limitations on the NA, as long as the working distance is sufficient to accommodate the additional components. The lateral resolution will still follow Abbe’s diffraction limit while the axial resolution will maintain as it mainly depends on the diffractive angle introduced by the grating.

We should note that the transmission grating inherently is wavelength dependent, which is inconvenient for simultaneous multi-color imaging and multi-photon implementation due to the dispersion. However, corrective optical design in the upstream optical path, for example, using refractive prism, can be implemented to compensate the dispersion. Albeit the limitations, the simple addition of a transmission grating in Meso-OPM provides a straightforward method to obtain 3D cellular resolution over a mesoscopic FOV, a significant capability for observing 4D dynamics in a large volume as demonstrated herein.

In summary, we circumvent the theoretical limitation of insufficient axial resolution in low NA objective lens by creating a high angle diffractive light sheet. By avoiding the trade-offs between FOV, axial resolution, and imaging speed, we demonstrate 4D cellular resolution over a FOV that is unattainable previously.

## Data availability

Data underlying the results presented in this paper are not publicly available at this time but may be obtained from the authors upon reasonable request.

## Supplemental document

See Supplement 1 for supporting content.

