## Supplement 1 for "Mesoscopic oblique plane microscopy (Meso-OPM) with a diffractive light sheet- enabling large-scale 4D cellular resolution imaging"

### 1. The characterization of the field of view (FOV) and resolution.

To evaluate the FOV and the resolution, red and green fluorescent beads with 1 μm diameter dispersed separately in agarose were imaged. The moving aluminum foil was synchronized to block the $-1_{st}$ and $0_{th}$ excitation during the imaging process (See *Methods* in Main text). 3350 images (2048 × 230 pixels) were acquired for each kind of fluorescent bead sample. As the excitation light sheet is oblique, the acquired images were oblique cross-sectional images. The real geometry of the acquired volume data was recovered by an affine transformation (See *Methods* in Main text). The imaging results for green and red fluorescent beads under excitation wavelengths of 488 nm and 561 nm are shown in Fig. S1 and S2, respectively. By converting the pixel dimension to the physical size with the calibrated magnifications, the FOV is calculated to be ~3.3 mm (X) × 5.4 mm (Y) × 0. 33 mm (Z). The FOV in the X direction is limited to 3.3 mm (2048×6.5/4) by the sensor size of the camera in our current setup, which is 2048 × 2448 pixels with pixel size of 6.5 μm. As the Y direction is the scanning direction, the FOV doesn't have such limitation as has in the X direction. Projections along the three axes of the volume datasets acquired for the green and red beads are shown in Fig. S1a-c) and Fig. S2 a-c, respectively. The brightness of the image is fairly uniform across the whole FOV except for slight vignetting at the edge of the FOV. Zoom-in views of the XY, XZ and the YZ projections are provided in Fig. S1d-f) and Fig. S2 d-f). The elongation of the beads along the axial direction is not significant so that the lateral profile of the beads tends to be very similar to that of the axial profile. This is a good indication that the axial resolution is very close to that of the lateral. Besides, the quality of the beads image is well maintained throughout the whole depth of ~330 μm except for minor deterioration due to the broadening of the excitation beam. To further examine the profile of individual beads, three representative beads in different depths are extracted. The projections of each bead are provided in Fig. S1h-j) and Fig. S2h-j) for green and red beads, respectively. The constant image quality of the beads in different depths further suggests that the resolution is well maintained in the accessible depth range. The comparison of the lateral and axial profiles between each bead further confirms that the axial resolution is significantly improved and comparable to that of the lateral. To quantify the resolution, we calculated the full width at half maximum (FWHM) of the line sections through maximum intensity projections of each bead. As for the 488 nm excitation, the average FWHM values across the FOV were 2.5 ± 1.1 ($x$), 3 ± 1.4 μm ($y$), 6 ± 1.8 μm ($z$) ($n$ = 50); as for that of 561 nm excitation, the average FWHM values were 2.9 ± 1 ($x$), 3.5 ± 1.5 μm ($y$), 6 ± 1.9 μm ($z$) ($n$ = 50). Short excitation wavelength has better lateral resolution, which is as excepted. But short excitation wavelength doesn't lead to better axial wavelength. This is probably because the angle of the diffractive light sheet under short excitation wavelength is smaller than that of the longer wavelength, which in turn cancels the axial resolution improvement under short wavelength. The measured axial resolution is

The axial resolution of an objective lens with NA (numerical aperture) of 0.3 is limited to ~11 μm under 488nm laser excitation. Therefore, the proposed method breaks this limitation and offers ~2-fold improvement in axial resolution.

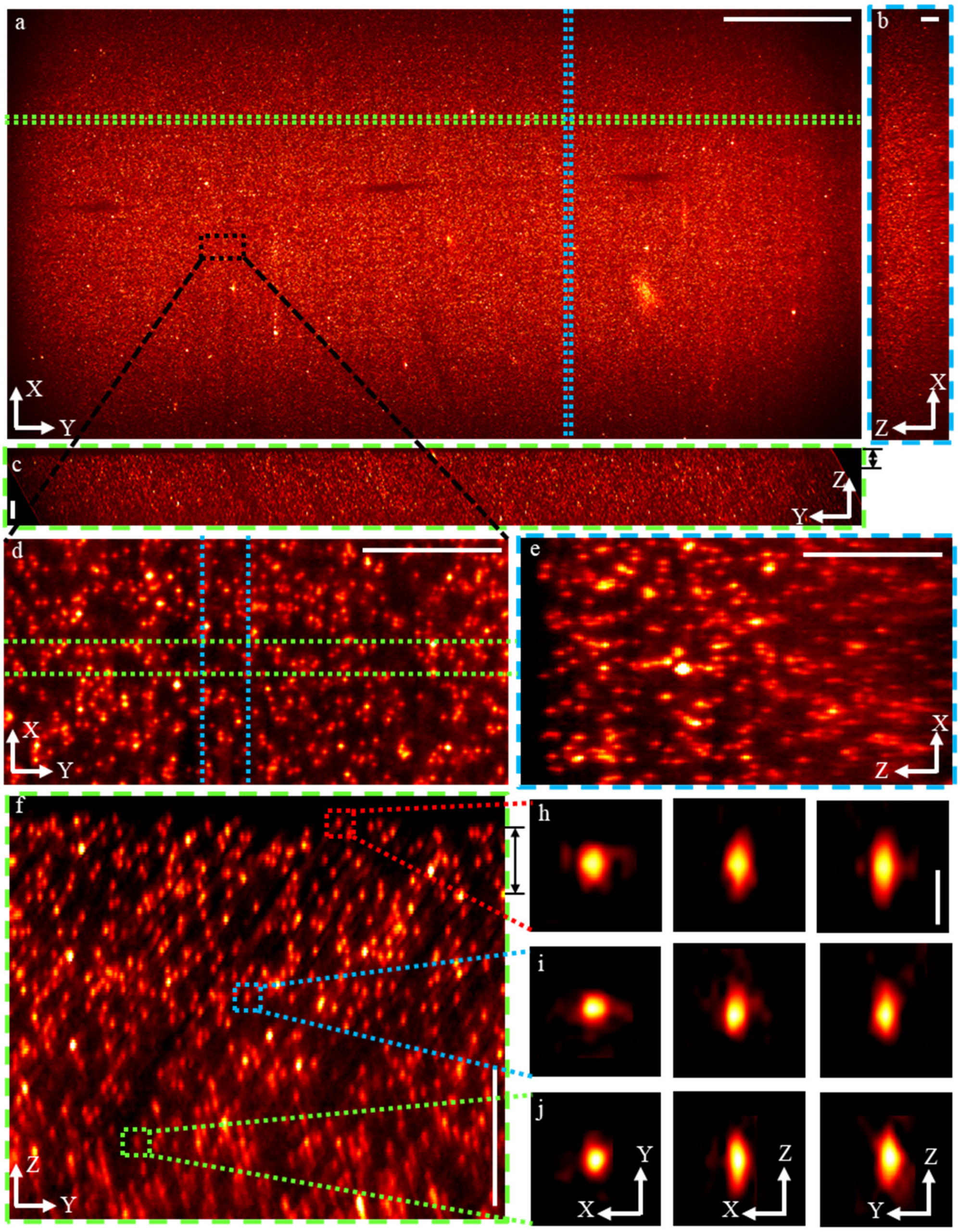


**Fig. S1. Resolution and FOV characterization under the blue laser (488 nm) excitation.** a) *En face* view generated by the MIP of the area between the black arrow in panel c). b) XZ cross-section generated by MIP of the area between the blue dash lines in panel a). c) YZ cross-section generated by MIP of the area between the green dash lines in panel a). d-f) Three zoom-in projections of the volume area that indicated by the black dashed rectangle in panel a). d) is the *en face* view generated by MIP of the depth layer indicated by the black arrows in panel f). e) and f) are the YZ and XZ cross-sections that are generated by MIP of the area indicated by the green dash lines and blue dash lines in panel d). The scale bar is 1000 μm for panel a), and the scale bar for the rest panels is 100 μm.

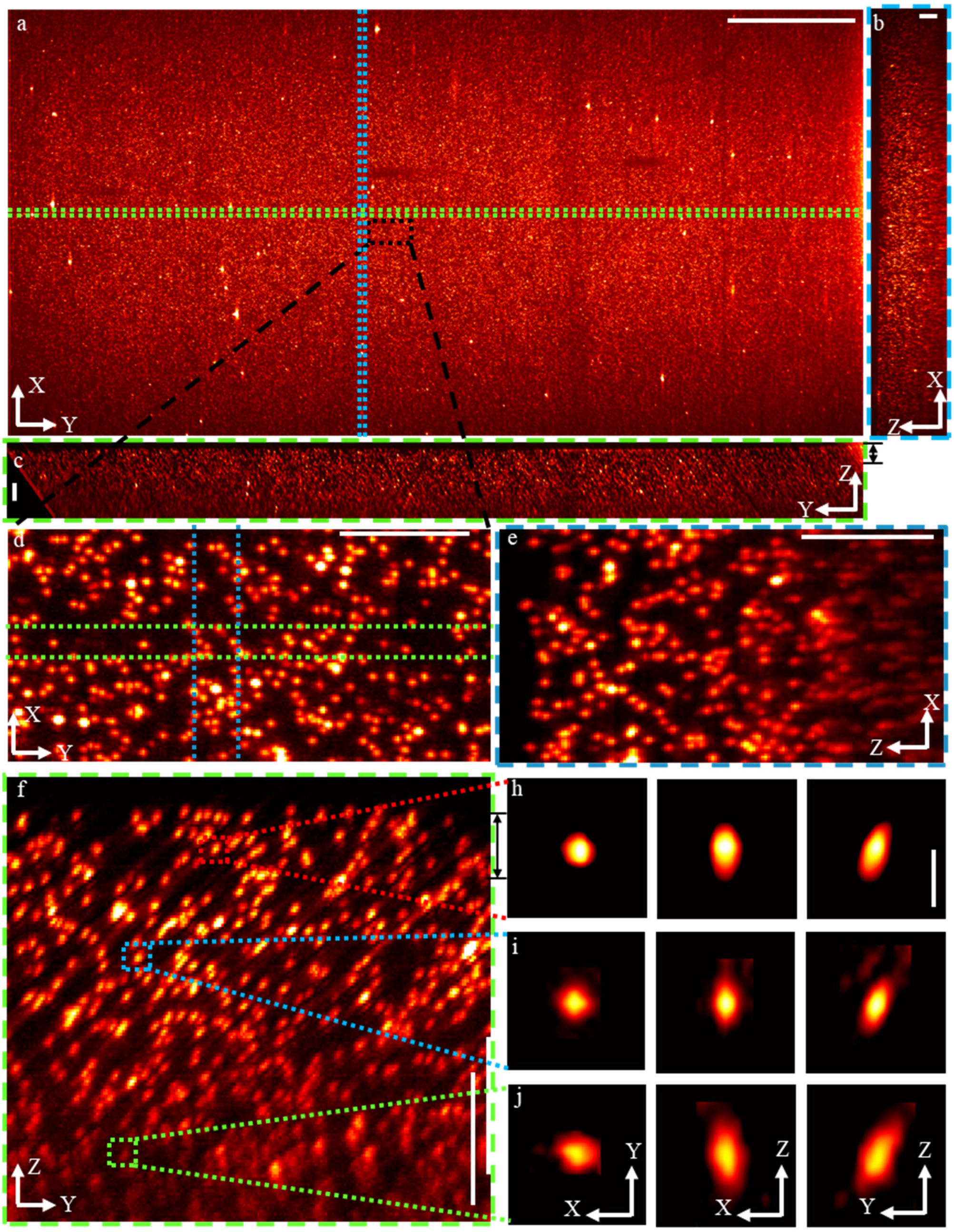


**Fig. S2. Resolution and FOV characterization under the green laser (561 nm) excitation.** a) *En face* view generated by the MIP of the area between the black arrow in panel c). b) XZ cross-section generated by MIP of the area between the blue dash lines in panel a). c) YZ cross-section generated by MIP of the area between the green dash lines in panel a). d-f) Three zoom-in projections of the volume area that indicated by the black dashed rectangle in panel a). d) is the *en face* view generated by MIP of the depth layer indicated by the black arrows in panel f). e) and f) are the YZ and XZ cross-sections that are generated by MIP of the area indicated by the green dash lines and blue dash lines in panel d). The scale bar is 1000 μm for panel a), and the scale bar for the rest panels is 100 μm.

### 2. The experiment setup

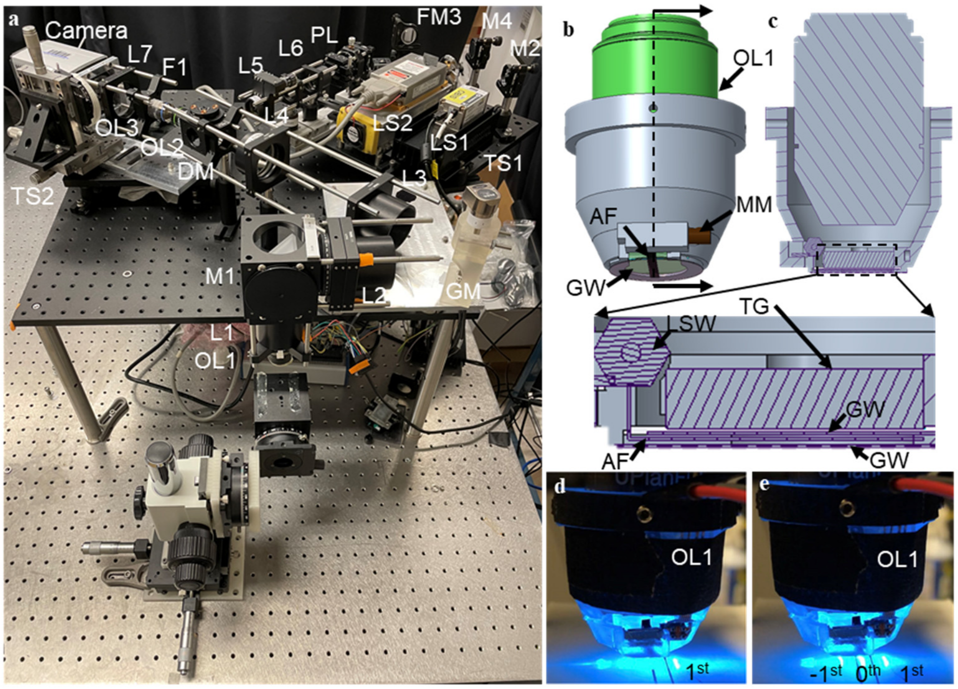


**Fig. S3. The real experiment setup and the 3D model of the primary objective lens.** a) The actual photograph of experiment setup; (L: lens; M: mirror; F: filter; PL: Powell lens; LS: light source; DM: dichroic mirror; GM: galvanometer mirror; TS: translation stage;) b) The 3D model of the assembly of the primary objective lens with the transmission grating (TG), aluminum foil (AF), micro-motor (MM), and glass window (GW); c) The cross-section of the assembly in panel (b) showing the alignment of the TG, GW, and AF. The AF is sandwiched between two GW and translated by the MM together with a micro linear stage (only the lead screw (LSW) of the linear stage can be seen); d-e) The real photograph of the assembly. As for panel d, the -1$^{st}$ and 0$^{th}$ orders are blocked by the aluminum foil so that only the 1$^{st}$ order can be used for excitation. As for panel e, the three diffraction orders (-1$^{st}$, 0$^{th}$, and the 1$^{st}$) are not blocked by the aluminum foil.

The actual photograph of the experimental setup is shown in Fig. S3a). Fig. S3b) is the 3D model of the assembly of the primary objective lens (OL1), TG, AF, GW, and MM. To illustrate the working principle of how the diffraction order of -1$^{st}$ and 0$^{th}$ are blocked, the internal structure of the assembly is provided in Fig. S3c). As shown in Fig. S3d), by moving the AF to a proper position, the -1$^{st}$ and 0$^{th}$ orders are blocked while the 1$^{st}$ is still can be used to illuminate the sample. On the contrary, without the engagement of the AF, all the three-diffraction orders can be observed. A video (supplement video 5) is also provided to demonstrate that the AF can be synchronized to block -1$^{st}$ and 0$^{th}$ orders of the moving light sheet.

### 3. The separation of light in different diffraction orders.

A ray tracing simulation shown in Fig. S4 is used to explain the separation of three diffraction orders (-1$^{st}$ 0$^{th}$, and the 1$^{st}$) generated by the transmission grating. The objective lens was designed according to the lens data of a public patent (10X objective lens) from Olympus. As the measured overall lateral magnification is ~4X, a paraxial lens with focal length of 72 mm is used to represent all the subsequent optics. As can be seen from Fig. S4a-c, the images of -1$^{st}$ and the 1$^{st}$ are defocused and have a distance of ~4 mm from the focused image of the 0$^{th}$ order. The 4 mm in the image space is equal to ~1 mm in the object space. Because the Rayleigh range of the light sheet is less than 1 mm along the Y direction (The diffraction direction is along the Y direction.), light comes from -1$^{st}$ and the 1$^{st}$ will not affect the image of the 0$^{th}$ order. The beads image shown in Fig. S1 and S2 also confirm that there are no artifacts caused by light from -1$^{st}$ and the 1$^{st}$. Please note that the light from the non-zero orders wasn't observed (or was very weak) even when utilizing the whole FOV of the camera which has a sensor size of ~13 mm × 16 mm. The reason might be due to the low diffraction efficiency when the incident angle is large.

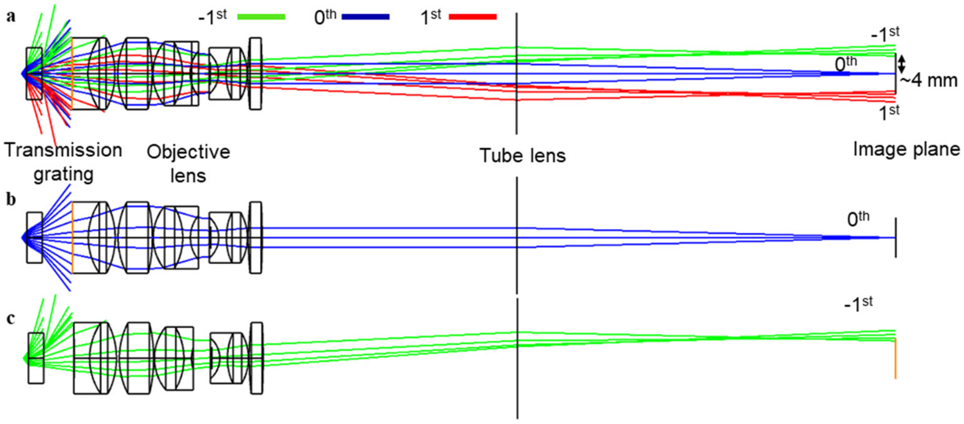


**Fig. S4. Ray tracing simulation of imaging with transmission grating.** a) Ray tracing simulation showing the images of three different diffraction orders are separated in the image plane. b) Ray-tracing simulations only showing the 0$^{th}$ order. c) Ray-tracing simulations only showing the -1$^{st}$ order.

### 4. The calculation of the light sheet parameters

Both the refraction and the diffraction can change the angle of the light sheet before it enters the sample. As shown in Fig. S5a, the light path of the light sheet is simplified for the calculation of the angle of the light sheet on the sample. $\theta_i$ (11°, see main text) is the incident angle of the excitation light. The light sheet is firstly bent by the diffraction of the transmission grating. The diffraction angle $\theta_m$ (~51° for 488nm and ~60° for 561nm) can be calculated according to the grating equation in the main text. Next, the angle of the light sheet will change due to refraction when light enters the water from the air. $\theta_r$ can be calculated by

$$\sin(\theta_r) / \sin(\theta_m) = n_{air} / n_{water} , \quad (S1)$$

where $n_{air}$ =1 is the refraction index of the air, and $n_{water}$ =1.33 is that of the water. As for 488 and 561nm, $\theta_r$ is calculated to be ~36° and 41°, respectively. Here we neglected the refraction by the thin cover glass due to smaller refractive contrast.

Figure S5b is used to calculate the NA of the focusing beam in water. $\Delta\theta_{air}$ and $\Delta\theta_{water}$ are half angles of light cone of the excitation light in air and water, respectively. The excitation beam has a radius of ~0.875 mm ($0.5 \times D_{beam} \times f_{L1} \times f_{L3} \times f_{L5} / f_{L2} / f_{L4} / f_{L6}$) when projected on the back aperture of the primary objective OL1. $D_{beam}$ = 0.7 mm is the beam diameter of the laser, $f_{L1}$ to $f_{L6}$ are the focal lengths of L1 to L6 (See methods section in main text). So, the NA ($NA_{ex\text{-}air}$) of the excitation beam in the air is ~0.875mm/18mm = 0.048 (18 mm is the focal length of OL1). Accordingly, the incident beam shown in Fig. S5b is within the range of ($\theta_m + \Delta\theta_{air}$) and ($\theta_m$ - $\Delta\theta_{air}$) in which $\Delta\theta_{Air}$ is the half angle of the light cone in air. $\Delta\theta_{air}$ can be calculated by taking the inverse sine of $NA_{ex\text{-}air}$. The range of refracted angle (($\theta_r + \Delta\theta_{water}$) to ($\theta_r$ - $\Delta\theta_{water}$)) in the water (Fig. S5b) can then be calculated by Eq. S1. $\Delta\theta_{water}$ is then calculated to be ~1.6° and ~1.4° for 488 nm and 561 nm, respectively. Given the half angle of the light cone, the beam waist ($\omega_0$) and Rayleigh ($Z_R$) range in water can be calculated as follows:

$$\omega_0 = M^2 \frac{\lambda}{\pi n_{water} \Delta\theta_{water}} \tag{S2}$$

$$Z_R = \frac{n_{water} \pi \omega_0^2}{\lambda} \tag{S3}$$

where $M^2$ (1.2 for 488 nm, 1.1 for 561 nm) is the beam quality factor, λ is the wavelength, $\Delta\theta_{water}$ needs to be converted to radians before calculation. As for 488 nm excitation, the beam waist and Rayleigh range in water are ~5 µm and ~214 µm, respectively. As for 561 nm excitation, the beam waist and Rayleigh range in water are ~6 µm and ~267 µm, respectively.

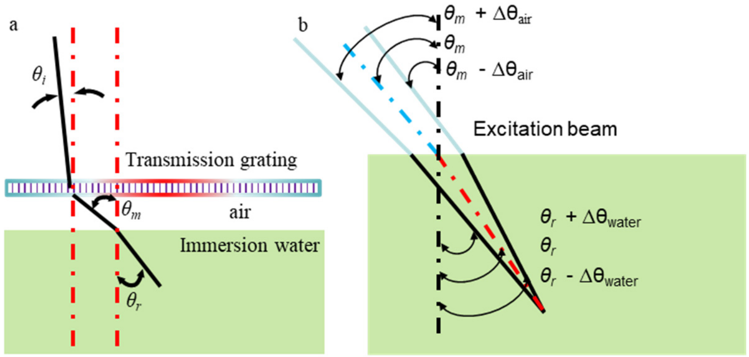


**Fig. S5**. The calculation of the angle and NA of the light sheet. a) The angle of the excitation light sheet is changed by both diffraction and refraction. b) The half angle of the light cone in air and water.

### 5. The numerical simulation for axial resolution

To evaluate the diffraction-limited resolution for the proposed method, a Fourier model for amplitude transfer function according to our previous publication[2] is established in Fig. S6a). The coordinate is reciprocal with the Cartesian coordinate (X, Y, Z) that is described in the main text (Main text: Fig. 1 and Fig. 2), and can be expressed with a wave vector $k$,

$$\vec{k} = \frac{2\pi}{\lambda} \hat{k}(k_x, k_y, k_z) \tag{S4}$$

where $\hat{k}$ is the directional vector with unit magnitude, λ is the wavelength (0.488 nm or 0.561 nm for different excitation wavelengths), $k_x$, $k_y$, and $k_z$ are the coordinates of the Fourier model. The blue circle $S_1$ and the orange circle $S_2$ represent the spatial frequency range for the excitation and the primary objective lens OL1 (See Fig. 2 in the main text), respectively. The black circle $S_3$ is the effective NA of primary objective lens because of the mismatched aperture between OL1 and OL2(See methods section in main text). The dash green circle $S_4$ shows the equivalent frequency range of the OL3 (See Fig. 2 in the main text) that is mapped on the back aperture of OL1. The overall spatial frequency range for the detection is illustrated by the overlapping area of $S_3$ and $S_4$. The definitions of $S_1$, $S_3$, and $S_4$ are then given as follows.

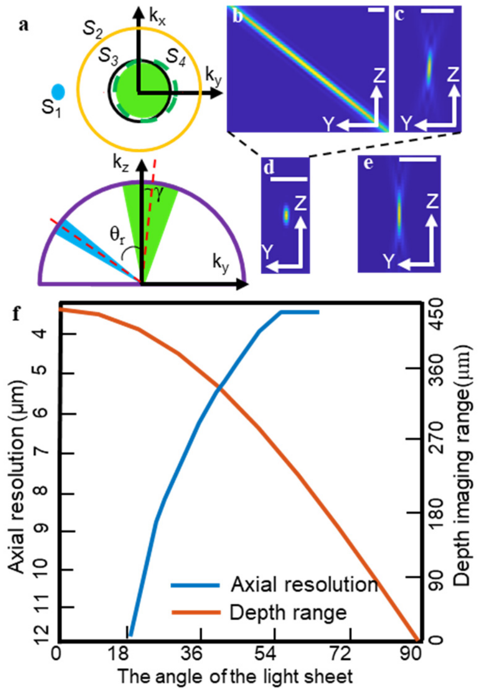


**Fig. S6. The calculation of the theoretical resolution and the depth detection range of the proposed method.** a) The 3D frequency support of the excitation and detection in 3D Fourier domain. b) and c) The Y-Z cross-sections of the excitation and detection PSFs. d) The Y-Z cross-sections of the combined PSF. e) The Y-Z cross-sections of the detection PSF under the full detection NA of 0.3. f) The The change of the axial resolution and the depth imaging range with that of the excitation angle $\theta_r$ under 488 nm excitation.

1) The definition of $S_1$

The effective NA of the excitation light in water can be calculated by $NA_{\text{ex-water}} = n_{\text{water}} \times \sin(\Delta\theta_{\text{water}})$ in which $\Delta\theta water$ (~1.6° for 488 nm and ~1.4° for 561nm) is obtained in the previous section. The special frequency range $S_1$ for the excitation light is identified as:

$$\left\{(kx,ky)\middle| \begin{array}{c}(k_x)^2+(k_y)^2+(k_z)^2=(2\pi/\lambda)^2 \\ \sqrt{k_x^2+(k_y+\sin\theta_r\times 2\pi/\lambda)^2+(k_z+\cos\theta_r\times 2\pi/\lambda)^2}\le(2\pi/\lambda)NA_{\text{ex-}water}\end{array}\right\}, \qquad \text{(S5)}$$

where $\theta_r$ is the oblique angle of the light sheet shown in Fig. S6a. $\theta_r$ is 36º for 488 nm laser and 41º for 561 nm laser (See previous section).

2) The definition of $S_3$

As the back aperture of the OL2 is overfilled (See methods section in the main text). The effective NA of OL1 is calculated as:

$$NA_{\text{eff-OL1}} = NA_{\text{OL1}} D_{OL2} / (M_{L1-L4} D_{OL1}) \approx 0.19\,, \qquad \text{(S6)}$$

where $NA_{\text{OL1}}$ = 0.3 is the NA of the OL1, $D_{OL1}$ = 10.8 mm and $D_{OL2}$ = 13.5 mm are the diameters of back aperture of OL1 and OL2, respectively. $M_{L1-L4}$ can be calculated as ($f_{L2}\times$

$f_{L4})/(f_{L1} \times f_{L3})$ = 2, which is the lateral magnification from of the relay lenses from L4 to L1. After obtaining $NA_{\text{eff-OL1}}$, $S_3$ can be written as:

$$\left\{(kx,ky)| \frac{(k_x)^2+(k_y)^2+(k_z)^2=(2\pi/\lambda)^2}{\sqrt{k_x^2+k_y^2+(k_z-2\pi/\lambda)^2}\leq(2\pi/\lambda)NA_{\text{eff-OL1}}}\right. . \tag{S7}$$

3) The definition of $S_4$

The equivalent collection NA of OL3 mapped on OL1 can be given as:

$$NA_{\text{eff-OL3}} = NA_{\text{eff-OL1}} NA_{\text{OL3}} / NA_{\text{OL2}}, \tag{S8}$$

where $NA_{OL2}$= 0.75, and $NA_{OL3}$= 0.75 is the NA of OL2 and OL3, respectively. $S_4$ can then be defined as:

$$\left\{(kx,ky)| \frac{(k_x)^2+(k_y)^2+(k_z)^2=(2\pi/\lambda)^2}{\sqrt{k_x{}^2+(k_y-\sin\gamma\times2\pi/\lambda)^2+(k_z-\cos\gamma\times2\pi/\lambda)^2}\leq(2\pi/\lambda)NA_{\text{eff}-OL3}\}}\right. , \tag{S9}$$

where the oblique angle $\gamma$ (Fig. S6a) of $S_4$ is due to the non-axial alignment OL2 and OL3. $\gamma$ can be estimated as follows:

$$\gamma = \varphi \times NA_{eff-OL1} / NA_{OL2}, \tag{S10}$$

where $\varphi$ is the angle between the optical axis of the OL2 and OL3 (19º for 488 nm excitation and 16º for 561 nm excitation, see methods section in main text).

The overall spatial frequency range for the detection is defined by the intersection area of $S_3$ and $S_4$, which can be obtained by Eq (S7) and (S9). The spatial frequency range for the excitation is given by Eq (S5). After the description of the 3D frequency support, the excitation and the detection amplitude point spread function (APSF) can be obtained by performing 3D Fourier transformation (Fig. S6, b-c). The intensity PSF is the squared magnitude of APSF. The combined system PSF can be calculated by Eq. 3 (See main text), which is a production of the excitation and the detection PSF.

The calculated theoretical resolutions are ~1.4 µm (X) × 1.6 µm (Y) × 6 µm (Z) for 488 nm excitation and ~1.5 µm (X) × 1.8 µm (Y) × 6 µm (Z) for 561nm excitation. The YZ cross-section of the system PSF is shown in Fig. S6d. The detection PSF from the full NA (0.3) of the OL1 has a FWHM of ~11 µm in the Z dimension (Fig. S6e), which can be calculated by the same method described above. According to the comparison in Fig. S6d-e, the proposed method can offer ~2 times better axial resolution than the diffraction-limited axial resolution of OL1. This also matches well with the results shown in Fig. S1 and S2.

To guide the choice of the line density of the transmission grating, we simulated axial resolution under different excitation angles of $\theta_r$. To simplify the calculation, we only change $\theta_r$ while keeping the detection $NA_{eff-OL3}$, $NA_{\text{ex-}water}$ and $\gamma$ *the* same as the above calculation. The change of axial resolution as a result of the different excitation angles is shown in Fig. S6f. There is a sharp increase in the axial resolution from 18º to 50º while the increment slows down when the angle is large than 50º. The axial resolution is not provided for the angle below 18º because the light collection issue of the remote focusing system will lead to axial resolution larger than 34 µm[1,3].

Due to the oblique excitation, the depth imaging range can be calculated by $2Z_R \times \cos\theta_r$, in which $2Z_R$ is twice the Rayleigh range (~214 µm for 488 nm excitation and 267 µm for 561µm nm excitation, see section 4). The change of the depth imaging range with that of the excitation angle $\theta_r$ is shown in (Fig. S6f). As for the current experiment, we chose a transmission grating with a line density of 0.83 μm/line to obtain oblique angle $\theta_r$ of 36° and 41° for 488 and 561 nm so that there is a balance between the axial resolution and the depth imaging range.

### 6. The full NA detection along the Y' axis

Figure S7 is used to illustrate the effective light collection angle of OL3 in the remote focusing system. The primary objective lens OL1 has an NA of 0.3. Due to the aperture mismatching, only partial light from OL1 can be collected by OL2. The effective NA of OL1 is calculated to be $NA_{\text{eff-OL1}}$ (0.19) by Eq. S6. The angle of the intermediate image plane is increased to 71° (under 488nm excitation) by the transmission grating and the demagnification design. The optical axis of OL3 is correspondingly rotated by 19° (90° - 71°) for correction. Given the angle of the intermediate image plane and the light cone of OL2(97°) and OL3(97°), the light collection angle of OL3 can be calculated to be 78°, resulting in light collection efficiency of >80% (78° /97°) from OL2 to OL3. Thus, the full NA detection of OL3 (under 488nm excitation) along the Y' axis can be calculated to be ~0.15 (0.19×78°/97°).

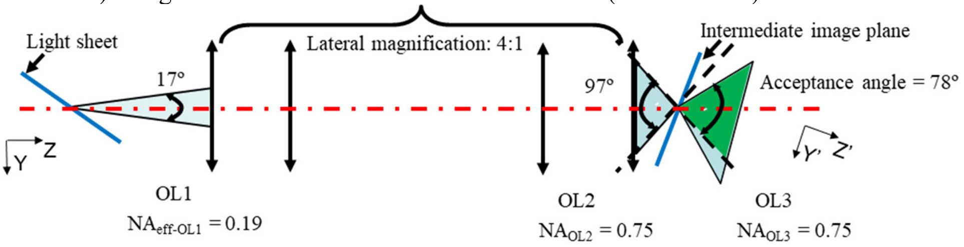


**Fig. S7. The light collection angle of the remote focusing system.**

### 7. The optical aberration associated with the transmission grating and immersion media

As the transmission grating has a thickness of ~ 2 mm and is placed underneath the objective lens, it may add additional optical aberration. Optical aberration may also increase as a result of using an air objective lens with immersion media. To evaluate the optical aberration associated with the transmission grating and the immersion media, a ray-tracing simulation shown in Fig. S8a) was designed in Zemax according to the real experimental setup described in the main text. Only the light from the 0$^{th}$ diffraction order of the transmission grating is utilized in the simulation. The thickness of immersion media is 1mm and has a refraction index that is close to that of water. The lens data for the two objective lenses are edited according to the public patents from Olympus. The achromatic lenses used as relay lenses (L1-L4) in the actual setup are replaced with the idea lens so that the comparison can reflect the optical aberration transmission that is caused by the grating and immersion media. As the back aperture of OL2 is overfilled, a reduced numerical aperture of ~0.19 (See Eq. S6) is used in the simulation. The wavelength in the simulation is 510 nm representing the center wavelength of emission spectrum in the green fluorescent protein, and the field point is ±1.65 mm covering the full FOV in the X-direction. Although the aberration is smaller without the use of transmission grating and the immersion media (Fig. S8b and c), the RMS in Fig. S8b) still suggests that the resolution can be maintained within ~1.5 μm when transmission grating and immersion media are used.

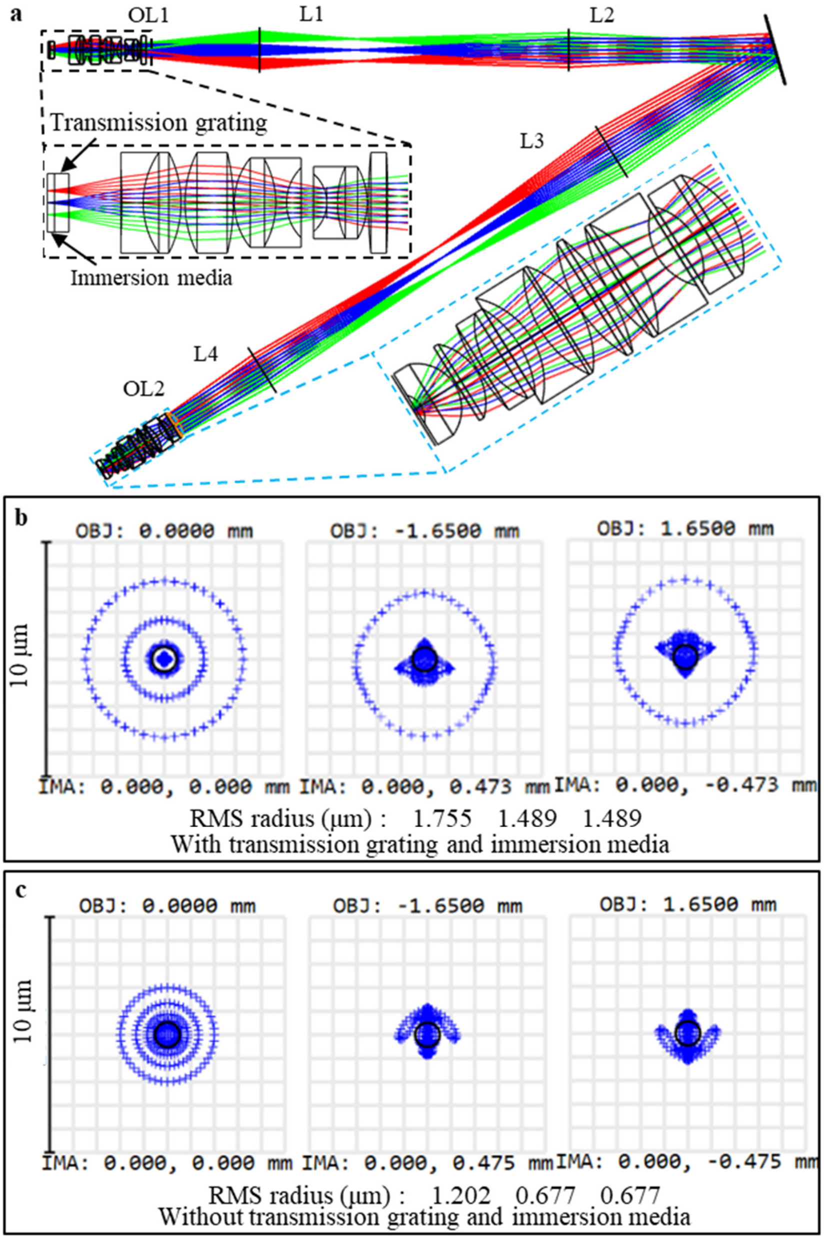


**Fig. S8. The evaluation of the optical aberration caused by the transmission grating and the immersion media.** a) The layout of the ray-tracing simulation. OL: objective lens; L: lens. b) The spot diagram with transmission grating and immersion media. c) The spot diagram without transmission grating and immersion media. RMS: root-mean-square. RMS is the root-mean-square value of the radius of all the rays.

### 8. Imaging parameters

| Figure Number | Camera frame rate (Hz) | Volume Rate | Camera FOV (pixels) | | | FOV (μm)$^2$ | | | Laser power (mW) | Excitation wavelength (nm) |
|---|---|---|---|---|---|---|---|---|---|---|
| | | | X | Y (Scans) | Z (Z') | X | Y | Z | | |
| Fig. 3 | 500 | 1.67 | 2048 | 300 | 275 (400) | 3.3 | 0.65 | 0.55 | 3.4 | 488 |
| Fig. 4 | 100 | 0.05 | 2048 | 2000 | 160 (230) | 3.3 | 5.4 | 0.33 | 2 | 561 |
| Fig. 5 | 250 | 2 | 2048 | 125 | 200 (290) | 3.3 | 0.5 | 0.4 | 3.4 | 488 |
| Fig. 6 | 625 | 5 | 2048 | 125 | 200 (290) | 3.3 | 0.5 | 0.4 | 3.4 | 488 |
| Fig. S1 | 100 | 0.03 | 2048 | 3350 | 160 (230) | 3.3 | 5.4 | 0.33 | 1 | 488 |
| Fig. S2 | 100 | 0.03 | 2048 | 3350 | 160 (230) | 3.3 | 5.4 | 0.33 | 1 | 561 |

**Table S1 Imaging parameters for each dataset shown**

The imaging parameters for all the datasets are summarized in table S1. As the pixel size will change after the affine transformation, both pixel sizes of Z′and Z are provided. The calibrated FOV in Z dimension is 0.33 mm. As for Fig 3, 5 and 6, the best image quality is within 0.33 mm in Z dimension although a larger volume is provided to show the whole sample.

### 9. The relationship between the FOV and working distance.

Figure S9 is used to illustrate how the FOV is changing with the working distance (WD). OL1 is the primary objective lens. WD is the distance from the focal plane of the OL1 to the grating surface. When the WD is longer the FOV becomes smaller accordingly.

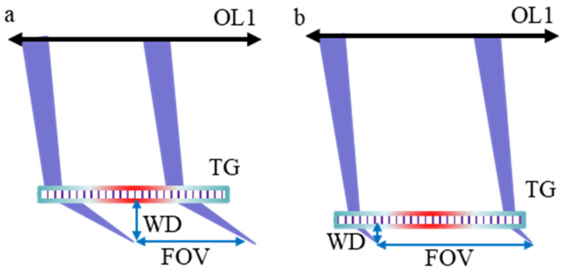


**Fig. S9**. The relationship between the FOV and working distance. a) Configuration with longer WD but smaller FOV. b) Configuration with shorter WD but larger FOV.

### 10. A comparison of the image acquired with Meso-OPM and confocal microscope.

To compare the image results with that of the confocal microscope, a zebrafish larva (4dpf, jGCaMP7s) was firstly imaged by Meso-OPM at 100 Hz per plane and scanned 300 planes at ~1.6 μm spacing. The results are shown in Fig. S10a-c. Figure S10d is *en face* z-projections of fish data acquired by a commercial confocal microscope (20X, NA/0.95). As the numerical aperture of the confocal microscope is much higher than that of the Meso-OPM, it offers a better lateral resolution. Individual neurons in the confocal image are much sharper than that of Meso-OPM, which is as expected. Nevertheless, Meso-OPM can still resolve dividual neurons, and similar anatomical organization of neurons can be observed in both images.

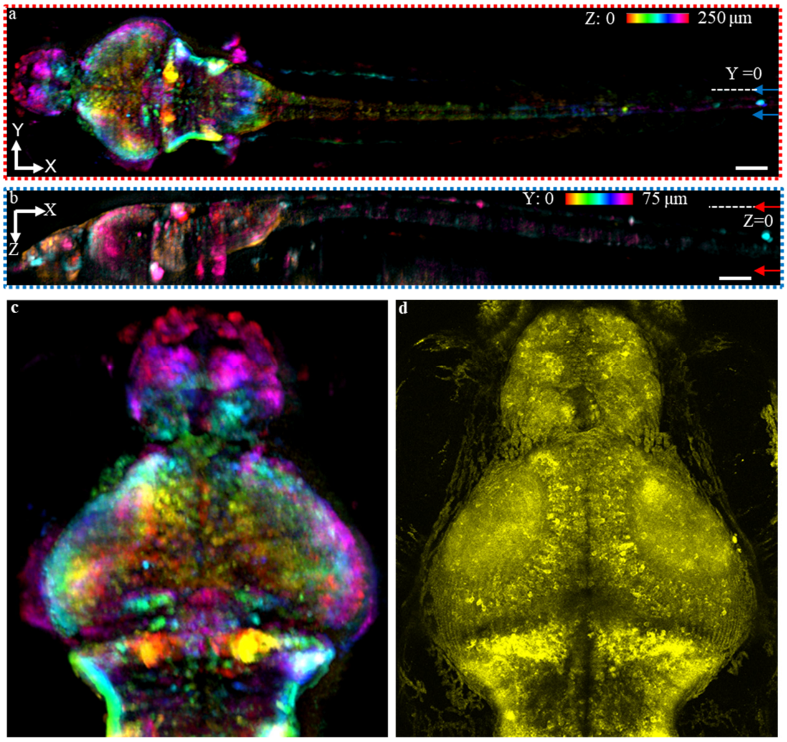


**Fig. S10. Image comparison between Meso-OPM (a-c) and confocal microscope (d).** (**a**) Color-coded MIP in XY plane over 250 µm along the Z dimension. The reference positions of the MIP are marked by arrows in panel b. (**b**) Color-coded MIP in XZ plane over 75 µm in the Y dimension. The reference positions of the MIP are marked by arrows in panel a. c) The zoomed *en face* z-projection imaged by Meso-OPM. e) The *en face* z-projections of a different larval fish imaged by confocal microscope.
